# A conditional protein diffusion model generates artificial programmable endonuclease sequences with enhanced activity

**DOI:** 10.1101/2023.08.10.552783

**Authors:** Bingxin Zhou, Lirong Zheng, Banghao Wu, Kai Yi, Bozitao Zhong, Yang Tan, Qian Liu, Pietro Liò, Liang Hong

## Abstract

Deep learning-based methods for generating functional proteins address the growing need for novel biocatalysts, allowing for precise tailoring of functionalities to meet specific requirements. This emergence leads to the creation of highly efficient and specialized proteins with wide-ranging applications in scientific, technological, and biomedical domains. This study establishes a pipeline for protein sequence generation with a conditional protein diffusion model, namely CPDiffusion, to deliver diverse sequences of proteins with enhanced functions. CPDiffusion accommodates protein-specific conditions, such as secondary structure and highly conserved amino acids (AAs). Without relying on extensive training data, CPDiffusion effectively captures highly conserved residues and sequence features for a specific protein family. We applied CPDiffusion to generate artificial sequences of Argonaute (Ago) proteins based on the backbone structures of wild-type (WT) *Kurthia massiliensis* Ago (KmAgo) and *Pyrococcus furiosus* Ago (PfAgo), which are complex multi-domain programmable endonucleases. The generated sequences deviate by up to nearly **400** AAs from their WT templates. Experimental tests demonstrated that the majority of generated proteins show unambiguous activity in DNA cleavage for both KmAgo and PfAgo, with many of them exhibiting superior activity as compared to the WT. These findings underscore CPDiffusion’s remarkable success rate to generate novel sequences for proteins of complex structures and functions in a single step with enhanced activity. This approach facilitates the design of enzymes with multi-domain molecular structures and intricate functions through *in silico* generation and screening, all accomplished without any supervision from labeled data.

## 1 Introduction

Deep learning-assisted functional protein design represents an innovative and promising approach that expedites and enhances the creation of novel proteins [1, 2]. By harnessing the power of deep learning, researchers are empowered to generate novel proteins with bespoke functions tailored to specific criteria [3–5]. The integration of deep learning in generating novel proteins offers opportunities to produce proteins with highly desired properties, such as higher stability, stronger binding affinity, and greater enzymatic activity. The exceptional capability of deep learning to generate and refine a diverse array of potential protein structures pushes the boundaries of protein engineering and discovery to new horizons. This expansion in scope offers opportunities for the creation of proteins with unique and valuable functionalities. Meanwhile, generating novel protein sequences of specific functions significantly augments researchers’ options, enabling the possibility of identifying protein candidates with superior activity and stability. Furthermore, the pool of diverse generated novel protein sequences enriches the library of the studied protein family over limited natural sequences. Such an augmentation not only supplements the resources available for the analysis and comprehension of proteins but also serves to provide an expanded group of protein templates for engineering toward enhanced functionality. At present, several prominent deep learning models have been applied in designing novel protein sequences of desired function [6–9]. One issue pertains to the extensive parameters of existing models, demanding a large amount of high-quality real-world data for training and fine-tuning, with substantial computational resources for deploying and inferencing. Meanwhile, extra efforts are required to experiment on the designed proteins by these models to discover a small subset of generated proteins that are soluble within buffers or display adequate bioactivity. Remarkably, the proteins tested in these researches tend to target relatively simple proteins in structure with a single domain, indicative of their simple functionality and obstacles generalizing to broader proteins or protein families. These collective challenges underscore the intricate nature of establishing new deep learning models to design novel sequences of proteins with multi-domain structures and complex functions.

As an emerging tool for generating diverse samples from complex distributions [10–14], denoising diffusion probabilistic models (DDPMs) stand out as natural and strong candidates for generating novel protein sequences. In the present task, identifying sequentially dissimilar proteins with desired structures or functions is of great significance. This requirement can be satisfied by the design philosophy of DDPMs, which assembles trainable neural network layers to progressively denoise artificial perturbations of defined noise distribution and reveal the original data distribution [9, 15–19]. Through iterative denoising steps aligned with specific objectives, DDPMs unravel the implicit mapping rules connecting a protein’s sequence and structure to its functions. Moreover, the denoising process can be conditioned on specific structural preferences or other characteristics of the protein of interest, guiding the output sampling distribution toward favorable directions. Consequently, a trained DDPM possesses the remarkable capability to generate diverse protein sequences conditioned by a set of desired properties tailored for the specific functionality.

This study practiced the DDPM methodology to generate novel sequences for programmable endonucleases called prokaryotic Argonaute (pAgo) proteins. pAgo protein is a class of endonucleases that play a crucial role in DNA interference in prokaryotes, and has gained significant attention in the field of biotechnology and bioengineering [20–25]. Their remarkable ability to target and cleave specific single-strand DNA/RNA sequences has led to important applications in diagnostics, which enables the design of molecular diagnostic assays for detecting and quantifying nucleic acid sequences associated with pathogens or cancer-related mutations [21, 23]. These assays offer improved early detection and precise treatment of diseases. Moreover, pAgo proteins exhibit high affinity to substrates and specific recognition of target sequences, making them valuable tools for imaging [26–28] and genes editing [29]. Mesophilic pAgo proteins are considered candidates for integration into isothermal nucleic acid-based detection and gene editing techniques [25, 30, 31]. However, their potential applicability is hindered by their low DNA cleavage activity. Thus, strategies are desired to enhance the enzymatic function of pAgo proteins at ambient conditions. In this context, we introduce a function-oriented design scheme using conditional protein diffusion (CPDiffusion) for sequence generation. CPDiffusion identifies a valid space of artificial pAgo sequences by learning implicit facilitation rules from a diverse set of data, including all different types of proteins with experimentally-solved structures and a small set of approximately 700 WT pAgo proteins. We generated two sets of long Ago proteins based on the backbones of *Kurthia massiliensis* Ago (KmAgo) and *Pyrococcus furiosus* Argonaute protein (PfAgo) [32, 33]. KmAgo is a mesophilic pAgo protein that can use both DNA and RNA as guide to cleave DNA and RNA, while PfAgo is a hyperthermophilic pAgo protein that can only use DNA as guide to cleave DNA and only functions at high temperatures in nature. Both of them have six structural domains and are nearly 800 amino acids in length. After an efficient training and screening procedure (Fig. 1), CPDiffusion delivers 27 novel artificial KmAgos (Km-APs) and 15 Pf-APs. They share 50 − 70% similarity in sequence identity compared to the template WT. The sequence identity of APs is less than 40% compared to other WT proteins from NCBI (except for the template). Unlike classic rational design methods, the entire process of model training and inference requires minimum expert guidance. Yet, when optimizing on a particular protein, CPDiffusion learns to automatically identify highly conserved regions while letting the remaining regions be highly variant.

**Fig. 1.**
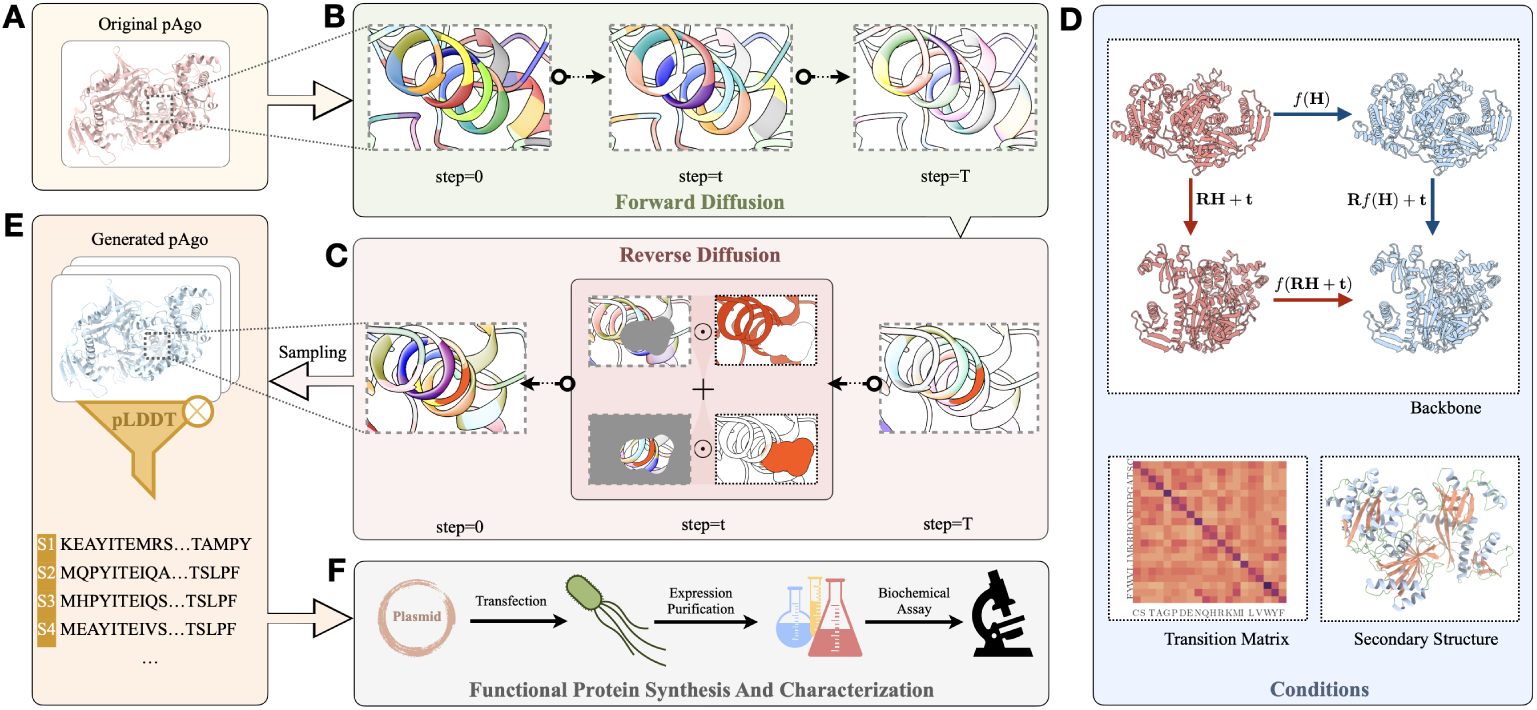
Workflow of CPDiffusion for designing novel mesophilic pAgo sequences. Arrows show the information flow among different components. (**a**) The WT prokaryotic Argonaute (pAgo) is processed to extract AA-level graph representation with molecule biochemical and topological properties (Section 1 in SI). (**b**) At the forward diffusion process, every AA type in the pAgo protein is corrupted in *T* steps following some transition probability matrix to gradually reach uniform distribution (Section 2.1 in SI). (**c**) The reverse diffusion process starts from randomly sampling each AA across the 20 AA types with a uniform distribution, following a progressive denoising process guided by (**d**) conditions, including the backbone template and secondary structure from the WT pAgo, and the transition matrix (Section 2.2 in SI). In particular, the propagation function *f* (·) is fitted by an equivariant graph convolutional layer that guarantees SE(3) equivariance with arbitrary rotation ***R*** and translation ***t***. (**e**) A set of protein sequences will be sampled from the learned distribution at denoising step 0, which is then screened based on the predicted pLDDT from AlphaFold2 [39] to confirm the candidates for (**f**) wet experimental synthesis, characterization, and evaluation.

We validated the enzymatic function and thermostability of CPDiffusion-designed pAgo proteins by biophysical and biochemical assays. Protein expression and purification experiments revealed successful expression and solubility of all 27 Km-APs. Significantly, 24 Km-APs displayed single-strand DNA (ssDNA) cleavage activity, with 20 of these sequences demonstrating superior ssDNA cleavage activity compared to the WT KmAgo. Meanwhile, all 15 Pf-APs were expressed and soluble in the buffer, displaying unambiguous ssDNA cleavage activity at 45 *^◦^*C, and the melting temperature shifts from 100 *^◦^*C to 50 *^◦^*C. Among these 15 Pf-APs, 6 Pf-APs (at 45 *^◦^*C) exhibited even greater activity than WT PfAgo at 95 *^◦^*C, with the enzymatic activity of the best Pf-AP being twice to that of the latter. The generation of novel programmable endonucleases by CPDiffusion with greatly enhanced activity at the ambient condition holds great practical value for easy-to-implement and high-throughput screening methods in gene editing and molecular diagnostics [25, 31, 34]. Moreover, the high success rate of CPDiffusion in generating novel sequences for large complex proteins, such as pAgo proteins with enzymatic function enhancement, represents a significant advancement in protein design and engineering for biomedical, biotechnological, and environmental applications.

## 2 Results

CPDiffusion trains a denoising diffusion model with 4 million learnable parameters from natural protein structure. The model assigns categorical noise distributions to AAs and learns their distribution via a conditional reverse diffusion. The overall construction of CPDiffusion are provided in the Methods section and Section 2-3 in SI. For baseline comparison, we trained a base model with ∼ 20, 000 crystal structures from **CATH 4.2** [35, 36] to learn sequence patterns from the conformation of protein structures. We validated the generative performance of the base model with the inverse folding task on several open test benchmarks spanning **CATH 4.2**, **TS50**, and **T500**. In the application of generating novel pAgo sequences, we trained CPDiffusion to learn the conservative patterns of Ago proteins on top of the general protein construction rules with WT proteins from **CATH 4.2**. We deliberately include a set of 693 WT proteins from the pAgo family to bias the model towards understanding pAgo proteins [37]. Note that the template protein used for generation was excluded from the full set of 694 pAgo dataset. We show in Section 5 in SI that training with additional pAgo family proteins enhances the model’s ability to generate sequences with desired AA composition patterns.

Compared to existing baseline methods, CPDiffusion generates reliable sequences with higher recovery rates and better sequence diversity (Section 4 in SI). The additional conditions, such as secondary structure and transition matrix, further attribute to an enhancement of CPDiffusion in recovering the distribution of protein sequences (Supplementary Fig. S3). Particularly, in the case of design sequences for pAgo proteins, CPDiffusion exhibits a significant advantage over baseline methods, such as ProteinMPNN. It does a notably better job of preserving biologically meaningful AA properties for proteins, such as polar AAs. This property is pivotal for protein stability and protein-nucleic acid interactions (Supplementary Fig. S4), yet ProteinMPNN misgenerated many polar AAs with opposite charges, potentially damaging the functionality of the generated proteins. Furthermore, the trained CPDiffusion successfully generated the catalytic tetrads in the PIWI domain, while ProteinMPNN fails (Supplementary Figs. S5-S6). Accurately generating catalytic tetrads is crucial for ensuring the functional integrity of the proteins in cleaving the target nucleic acids, and mis-generation of any AA within the tetrads designed for the studied pAgo proteins would directly result in failure of their function [38].

### Screening Workflow for Generative Ago Sequences

The trained CPDiffusion was applied to generate 100 pAgo sequences guided by the KmAgo template with *<* 70% sequence identity to the template. An initial structure-based screening is next processed to reduce the experimental workload. Given the structural conservation of pAgo proteins, we adopted AlphaFold2 [39] to assess the quality of the new sequences by their structural prediction confidence with respect to the template KmAgo. Initially, the overall pLDDT (predicted Local Distance Difference Test) scores of the generated sequences were compared against each other, where the generated samples with their overall pLDDT scores lower than one standard deviation below the average performance among 100 samples were eliminated (Fig. 2a, top panel). Next, the residue-level discrepancies of generated samples and WT KmAgo were measured by *σ*(ΔpLDDT) (Fig. 2a, middle panel) and count(|ΔpLDDT| *>* 10) (Fig. 2a, bottom panel), where ΔpLDDT = pLDDT_AP_*_i_* − pLDDT_WT_ describes perresidue pLDDT difference between the generated and WT KmAgo sequences. The former calculates the volatility of per-residue pLDDT difference between APs and the WT. As we desire sequences with a smaller inconsistency with their WT template, those with *σ*(ΔpLDDT) exceeding one standard deviation above the average inconsistency were removed. The last criterion disposes sequences of more than 93 AA positions that have ΔpLDDT larger than 10. Similarly, the threshold 93 was taken from one standard deviation above the average.

**Fig. 2.**
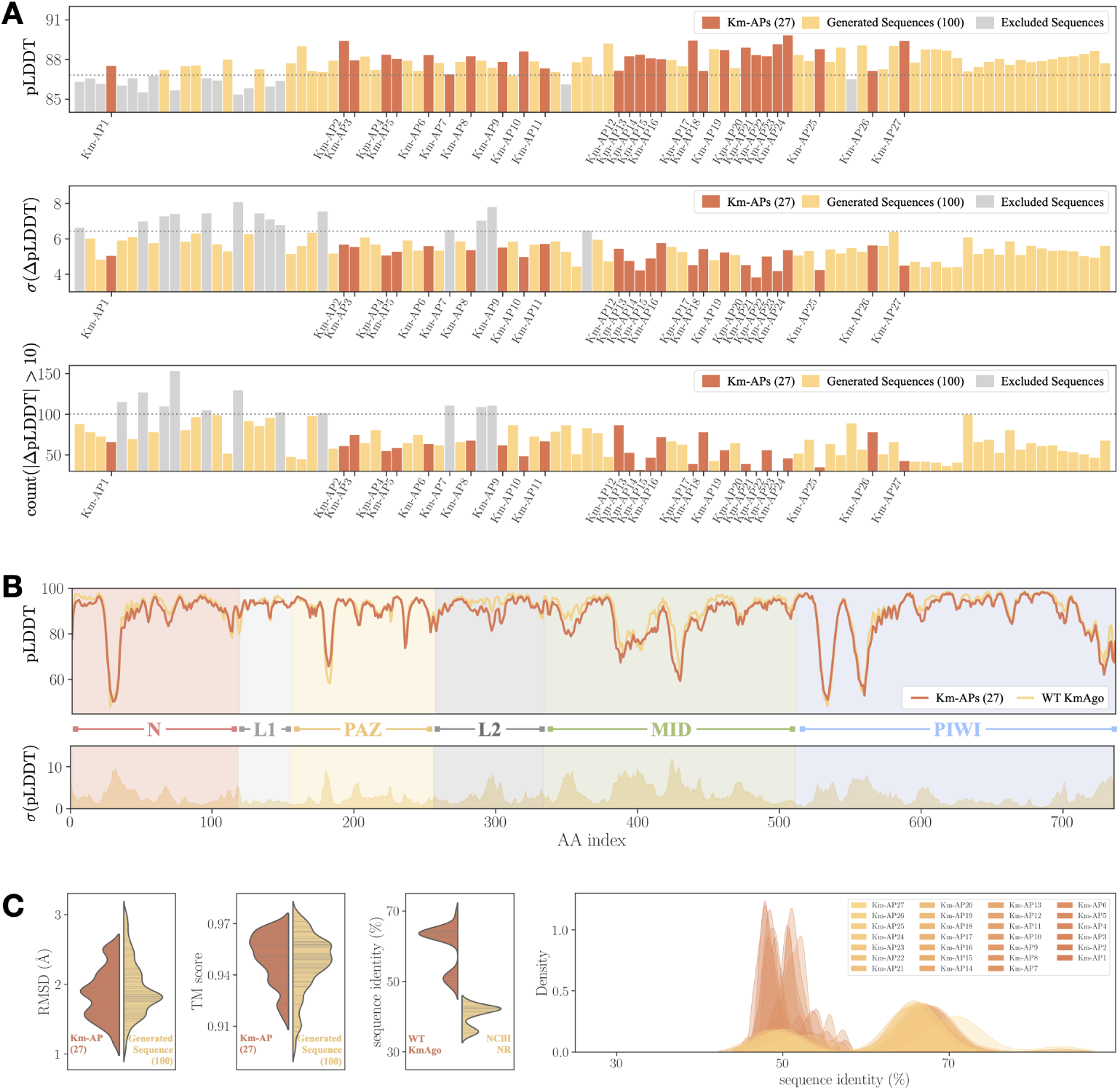
Initial screening and sequence-based analysis for CPDiffusion-generated artificial KmAgo proteins. (**a**) The three steps for the initial screening of the generated 100 sequences (yellow). The excluded sequences at each step are colored in gray, and the final 27 Km-APs are highlighted in red. The filtering thresholds are displayed as dotted horizontal lines. The three panels from top to bottom visualize the protein-wise average pLDDT of generated KmAgo sequences (top), the average volatility of residue-level discrepancies between WT KmAgo and the generated KmAgo sequences (middle), and the count of AA sites with large residue-level pLDDT discrepancies (bottom). (**b**) Comparison of residue-wise pLDDT for WT KmAgo and the average Km-APs (upper panel, in red). The standard deviation of the pLDDT difference between WT KmAgo and Km-APs is displayed in the bottom panel. (**c**) Summary of the structure (the left two subplots) and sequence variance (the right two subplots) for generated KmAgo sequences. The RMSD (first subplot) and TM score (second subplot) of the 27 Km-APs (in red) and 100 generated sequences (in yellow) with respect to WT KmAgo indicate an overall consistency of the AlphaFold2-predicted structure between generated sequences and WT KmAgo, with a few outliers in the group of 100 generated sequences. For the 27 Km-APs, their sequence identity (third subplot) is between 50 − 70% to WT KmAgo (red) and between 30 − 45% to all WT proteins in the NCBI NR database (yellow). Moreover, their pairwise sequence identities (fourth subplot) are below 40 − 80%, with the majority falling near 50%.

The three initial screening criteria outlined above led to the exclusion of ∼ 30 of 100 generated sequences. A more refined analysis is pursued subsequently by comparing the pLDDT scores of generated sequences and WT KmAgo at the residue level. Eventually, we selected 27 Km-APs (the sequences are listed in Supplementary Tables S5-S13) that exhibit high consistency in local structures (predicted by AlphaFold2) with WT KmAgo (Fig. 2b and Supplementary Figs. S8-S34). In contrast, example pLDDT curves of excluded APs are shown in Supplementary Figs. S35-S40 for comparison, where they all exhibit some local inconsistency. Additionally, the selected Km-APs underwent structural alignment with the WT KmAgo to confirm their structural similarity (Supplementary Fig. S42). The structural discrepancy was also quantified using Root Mean Square Deviation (RMSD) and Template Modeling (TM) scores (Supplementary Fig. S41), where the RMSD of all 27 selected proteins with WT KmAgo is below 3 Å, falling within the resolution range typically observed in X-ray crystallography of pAgo protein [38, 40, 41]. Meanwhile, TM scores of all Km-APs are above 0.9, suggesting an identical 3D structure of the generated and WT KmAgo. The high structural consistency is not only observed in the 27 chosen Km-APs but also for all CPDiffusion-generated sequences. As summarized in Fig. 2c, both Km-APs (red) and CPDiffusion-generated raw sequences (yellow) exhibit low RMSD and high TM scores. In other words, the screening procedure selects the most confident protein sequences for testing rather than predicting inactive samples to filter out. In addition to high structural consistency, the generated protein sequences also demonstrate high sequential diversity. As displayed in Fig. 2c (the right two panels), the 27 Km-APs are 50 − 70% similar to WT KmAgo and 30 − 40% similar to all WT proteins included in the NCBI NR database [42]. Moreover, the pairwise identity among these generated Km-APs ranges between 40 − 70%. More detailed statistics can be found in Supplementary Figs. S44-S47.

The overall procedure of generating and screening remains the same for PfAgo. In total, we selected 15 Pf-APs from 50 generated sequences based on the WT PfAgo template (See Section 6.4 in SI). Similar to Km-APs, the 15 Pf-APs also exhibit high consistency in structure construction. All TM scores are above 0.97, and all RMSD scores are below 1 Å (Supplementary Fig. S83). The sequence identities are ∼ 60% compared to WT PfAgo and ∼ 40% compared to non-PfAgo proteins (Supplementary Figs. S86-S88). The sequences for experimental examination are listed in Supplementary Tables S19-S23.

### Experimental Assessment of Artificial Proteins on Solubility, Activity, and Thermostability

The previous screening procedure returned 27 Km-APs that exhibit consistent structures with WT KmAgo (Fig. 3a and b) and satisfying diversity at the sequence level. Wet-lab experiments were then proceeded to evaluate the expression, single-strand DNA (ssDNA) cleavage activity, and thermostability of these Km-APs. Protein expression was assessed using a polyG linker fused with green fluorescence protein (GFP) for each Km-AP. Under buffer conditions, all 27 Km-APs displayed notable green fluorescence signals (Fig. 3c), indicating successful expression and solubility. We conducted synchrotron small-angle X-ray scattering (SAXS) to test the status of Km-APs in solution. We calculated the radius of gyration (R*_g_*) of proteins in buffer using Guinier analysis and the pair distribution function (P(r)). As shown in Supplementary Fig. S50 and Supplementary Table S14, both analyses demonstrate that the R*_g_* of Km-APs is the same as Km-WT, indicating that the Km-APs remain as monomer in buffer. The foldability of Km-APs was investigated by comparing the circular dichroism (CD) signal and SAXS to that of Km-WT, with the results in Supplementary Fig. S51 and Supplementary Table. S14 indicating correct folding in the buffer. Subsequently, we examined the ssDNA cleavage activity of these Km-APs. Fig. 3d presents the cleavage process for pAgo proteins, where red fluorescence indicates effective cleavage of the target DNA (tDNA). Remarkably, 24 Km-APs demonstrated ssDNA cleavage activity at 37*^◦^*C (Fig. 3e, the protein quantification of activity experiments is shown in Supplementary Fig. S58, the cleavage assay achieved by protein expressed *in vitro* shown in Supplementary Fig. S52), and 20 Km-APs exhibited enhanced DNA cleavage activity compared to WT KmAgo. The best-performing protein (Km-AP23) demonstrated nearly 9 times the ssDNA cleavage activity of the WT. We conducted additional experiments to further test the cleavage activity of the Km-APs with significantly enhanced functions. In these two experiments, the ratios of protein:guide:target were as 5:2:2 and 3:2:2. The results shown in Supplementary Fig. S59 indicate that the Km-APs exhibit enhanced cleavage activity under various protein:guide:target ratios. These results further demonstrate that CPDiffusion can generate Km-APs with enhanced functions. To further validate the enhancement in functionality of Km-AP23, we quantified the biofunction of Km-WT, Km-AP23, and Km-AP9 (reduced function) by using the Michaelis-Menten kinetic model. As shown in Supplementary Fig. S60 and Supplementary Table S15, we found the K*_M_* value for Km-AP23 is decreased and that for Km-AP9 is increased compared with Km-WT, which indicates that the affinity of Km-AP23 for substrates is enhanced, while that of Km-AP9 is reduced. Additionally, the k*_cat_* value for Km-AP23 is increased and that for Km-AP9 is decreased compared with Km-WT, which suggests that the cleavage efficiency of Km-AP23 is enhanced, while that of Km-AP9 is reduced. These results further demonstrate that Km-AP23 has higher activity than Km-WT. We also compared Km-AP23 with other well-established WT pAgo proteins (Supplementary Fig. S61) and conducted cleavage assays on different guide and target DNA sequences from various diseases and viruses (Supplementary Fig. S62). Km-AP23 consistently exhibited higher cleavage activity than other mesophilic pAgo proteins across different DNA sequences. To provide more insights into the performance of Km-AP23, we performed the electrophoretic mobility shift assay (EMSA) to analyze the binding of Km-AP23 to gDNA. The results, presented in Supplementary Fig. S53, show that compared with Km-WT, the formation of the protein-gDNA binary complex is increased in Km-AP23 at various protein concentrations. This observation demonstrates enhanced gDNA binding by Km-AP23, which can then provide more templates for enhanced binding to tDNA. We also employed the fluorescence polarization assay to study its binding affinity to gDNA and tDNA. The results in Supplementary Fig. S54 showed that the dissociate constant (K*_d_*) of Km-AP23 is lower than that of WT KmAgo, indicating an increased binding affinity of Km-AP23 to gDNA and tDNA. Furthermore, we tested the nucleic acid preference of Km-AP23 (Supplementary Fig. S63). Km-AP23 displayed enhanced cleavage activity on both DNA and RNA when utilizing ssDNA as a guide. However, it demonstrated either comparable or reduced cleavage activity on DNA and RNA when employing ssRNA as a guide, indicating that Km-AP23 is more likely to use ssDNA as a guide than ssRNA [32, 43]. We calculated the binding free energy of gRNA/tDNA for the entire structure of Km-WT and Km-AP23 using molecular dynamics (MD) simulations (the structures of the protein-gRNA-tDNA complex are obtained from AlphaFold3). As shown in Supplementary Fig. S64, the binding free energy of gRNA/tDNA for Km-AP23 is higher than that for Km-WT, which suggests that Km-AP23 may not form a stable protein/gRNA/tDNA structure to perform cleavage. Additionally, we analyzed the distribution of mutation sites in Km-AP23. As shown in Supplementary Fig. S102, the mutations are more frequently found on the protein surface. The thermostability of Km-APs was evaluated by incubating them at 42*^◦^*C for 2 and 5 minutes, followed by assays for ssDNA cleavage activity, which were then normalized to their respective activities at 37*^◦^*C. Fig. 3f shows that 10 of 27 Km-APs exhibited greater thermostability than WT KmAgo. Particularly noteworthy were some Km-APs, such as Km-AP5 and Km-AP14 (Fig. 3f), which showed concurrent enhancements in both activity and thermostability.

**Fig. 3.**
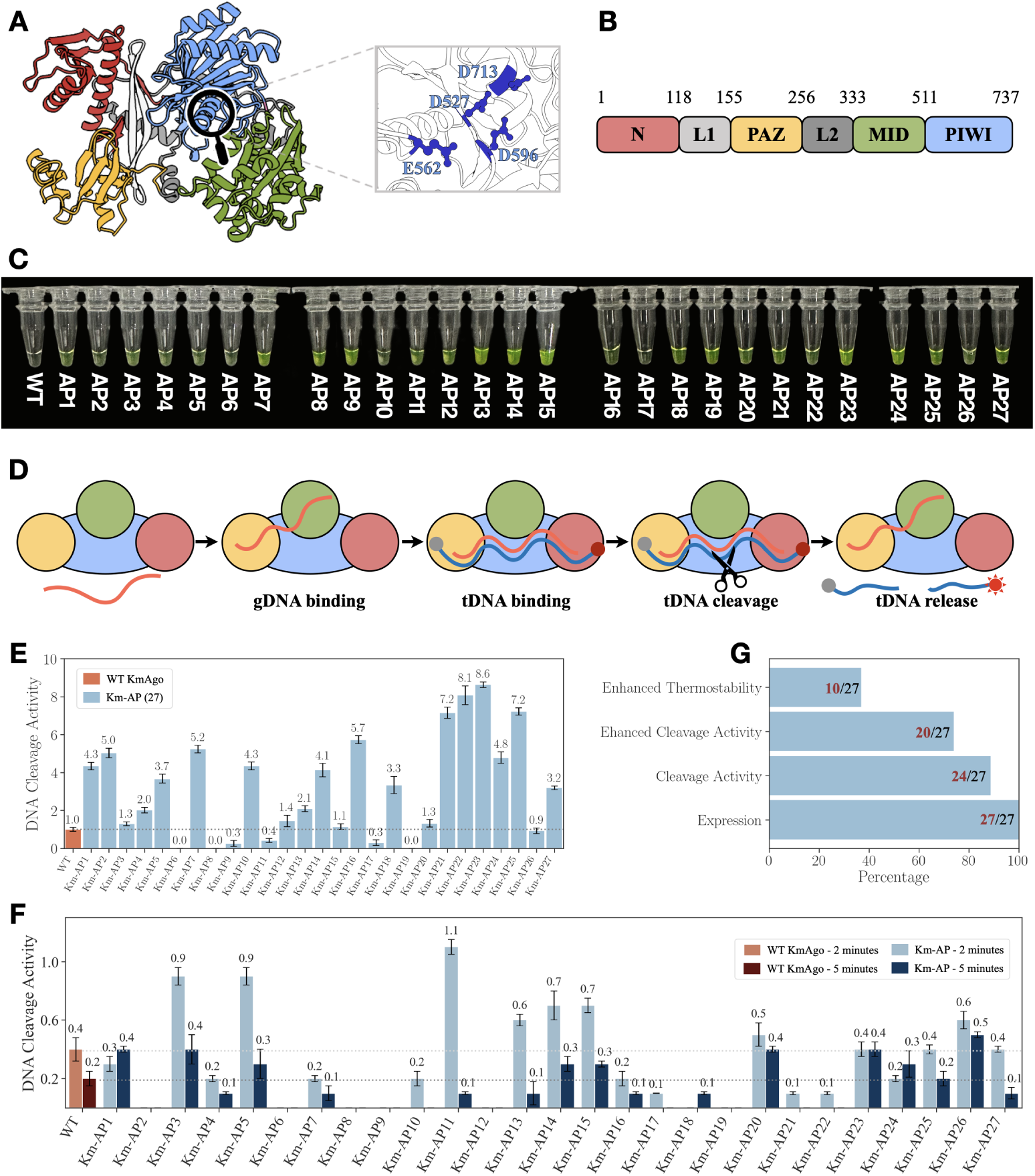
Synthesis and characterization of CPDiffusion-generated artificial KmAgos. (**a**) Left panel: Representation of the domain architectures of WT KmAgo (predicted by AlphaFold2). The N domain, Linker1, PAZ domain, Linker2, MID domain, and PIWI domain are colored in red, light gray, yellow, green, gray, and blue, respectively. Right panel: Catalytic sites of WT KmAgo. (**b**) Schematic diagram of the domain organization of WT KmAgo. (**c**) 27 Km-APs exhibited soluble expression, visualized by linking GFP to Km-APs. (**d**) Schematic diagram of the DNA-catalytic cycle of KmAgo. (**e**) Cleavage activity of WT KmAgo and 27 Km-APs at 37*^◦^*C on ssDNA. Proteins are loaded with ssDNA guide and then incubated with ssDNA target in a 2 : 1 : 1 molar ratio (protein:guide:target). These activities are defined by the slope of the time dependence of fluorescence intensity [44]. Results from two independent experiments were quantified, with the associated standard deviations represented by error bars. (**f**) Cleavage activity of WT KmAgo and 27 Km-APs after being incubated at 42*^◦^*C for 2 minutes and 5 minutes before conducting ssDNA cleavage assays to examine the thermostability enhancement of the Km-APs in comparison to WT KmAgo. (**g**) Summary performance of the 27 experimented Km-APs, where all of them can be expressed, with 24 exhibiting single-strand DNA cleavage activity, 20 surpassing the WT’s cleavage activity, and 10 showing enhanced thermostability.

For Pf-APs, we selected 15 Pf-APs for wet experiments validation. Although the structure of KmAgo and PfAgo is considerably conservative (Fig. 4a and b), with a sequence identity of 25.42%, the two proteins diverge at different evolutionary branches (Fig. 5a), and they have distinct active temperatures and nucleic acid preferences [32, 33, 44]. The SDS-PAGE and SAXS experiments demonstrated that all Pf-APs can be purified and remain as monomers in the buffer, respectively (Supplementary Fig. S90 and Supplementary Table S24). The overall packing of Pf-APs was investigated by comparing the SAXS to that of Pf-WT, with the result in Supplementary Fig. S91 indicating correct folding in the buffer. The ssDNA cleavage assay was performed on these Pf-APs, and remarkably, all 15 Pf-APs demonstrated ssDNA cleavage activity (Fig. 4c, the protein quantification of activity experiments is shown in Supplementary Fig. S93, the cleavage activity experiments under the different ratios of protein:guide:target are shown in Supplementary Fig. S94). To comprehensively assess their performance, we used PfAgo’s ssDNA cleavage activity at both 45 *^◦^*C and 95 *^◦^*C, and KmAgo’s ssDNA cleavage activity at 45 *^◦^*C as references. All Pf-APs exhibited enhanced cleavage activity compared to WT PfAgo at 45 *^◦^*C. Furthermore, 11 Pf-APs showed enhanced cleavage activity compared to WT KmAgo at 45 *^◦^*C, and 6 Pf-APs (at 45 *^◦^*C) were even more active than WT PfAgo at 95 *^◦^*C (Fig. 4e). A notable observation is that the melting temperature (∼ 50*^◦^*C) of Pf-APs is lower compared to WT PfAgo (∼ 100 *^◦^*C) (Fig. 4d), and the activity of Pf-APs is enhanced at moderate temperatures. These results can be attributed to the fact that our training dataset primarily consists of pAgo proteins from mesophilic prokaryotes (Supplementary Fig. S101). Thus, we can conclude that Pf-APs generated by CPDiffusion maintain the functional feature from the template WT while their thermostability, associated with the overall AA packing, is inherited from the training dataset. This example might pave a new route for future applications to engineer a hyperthermophilic protein to work at the mesophilic condition.

**Fig. 4.**
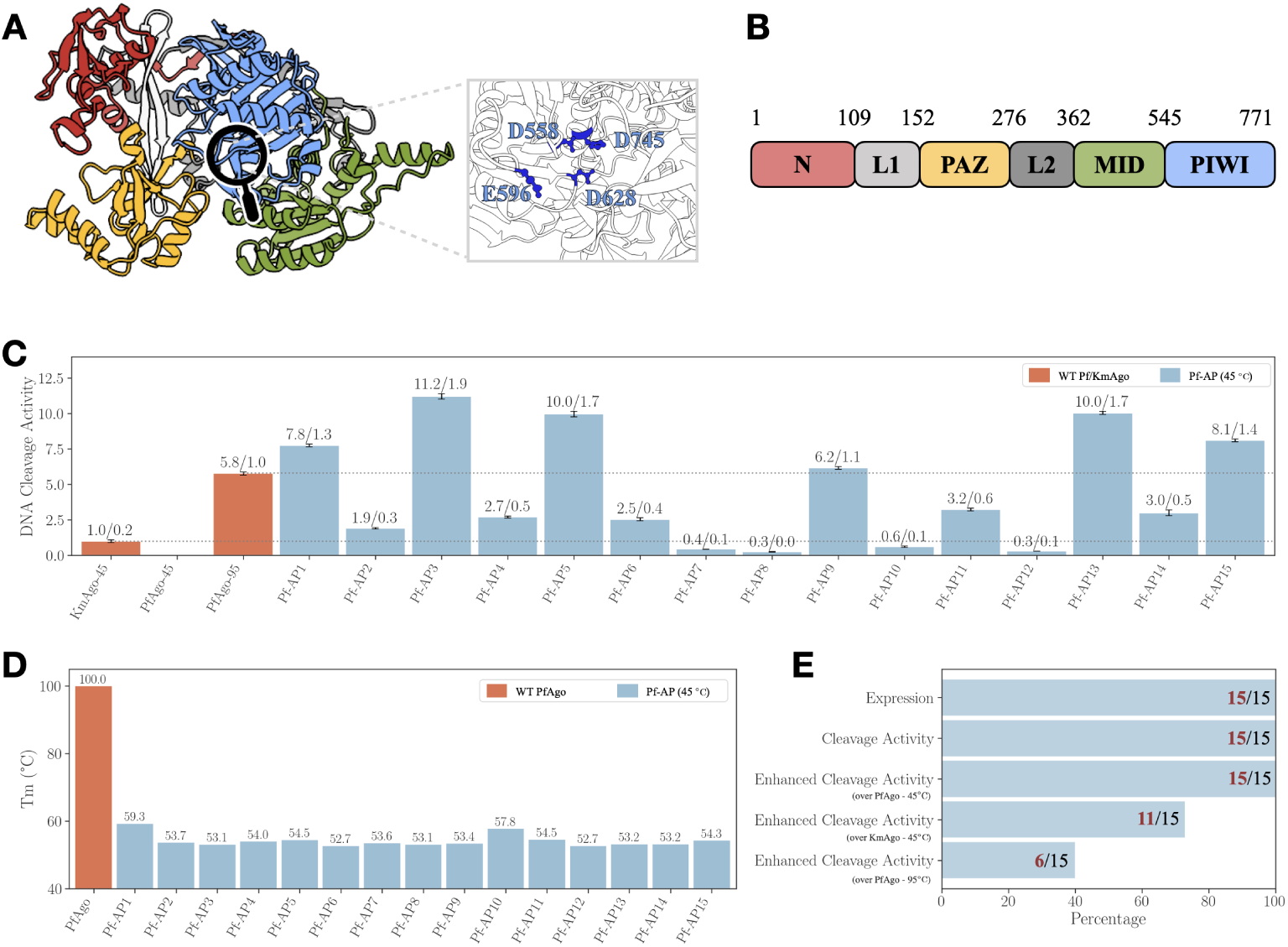
Synthesis and characterization of CPDiffusion-generated artificial PfAgos. (**a**) Left panel: Representation of the domain architectures of WT PfAgo (predicted by AlphaFold2). The N domain, Linker1, PAZ domain, Linker2, MID domain, and PIWI domain are colored in red, light gray, yellow, green, gray, and blue, respectively. Right panel: Catalytic sites of WT PfAgo. (**b**) Schematic diagram of the domain organization of WT PfAgo. (**c**) Cleavage activity of 15 Pf-APs at 45*^◦^*C. Proteins are loaded with ssDNA guide and then incubated with ssDNA target in a 2 : 1 : 1 molar ratio (protein:guide:target). The activities are defined by the slope of the time dependence of fluorescence intensity [44, 52]. Results from two independent experiments were quantified, with the associated standard deviations represented by error bars. (**d**) Melting Temperature (Tm) of WT PfAgo and Pf-APs. Tm of Pf-APs is determined by DSF spectra, and Tm of WT PfAgo is retrieved from [44]. (**e**) Out of the 15 experimented Pf-APs, all of them can be expressed, exhibiting single-strand DNA cleavage activity, and surpassing the WT PfAgo’s cleavage activity at 45 *^◦^*C. Additionally, 6 and 11 Pf-APs surpass the WT PfAgo’s cleavage activity at 95 *^◦^*C and exceed the WT KmAgo’s cleavage activity at 45 *^◦^*C, respectively.

**Fig. 5.**
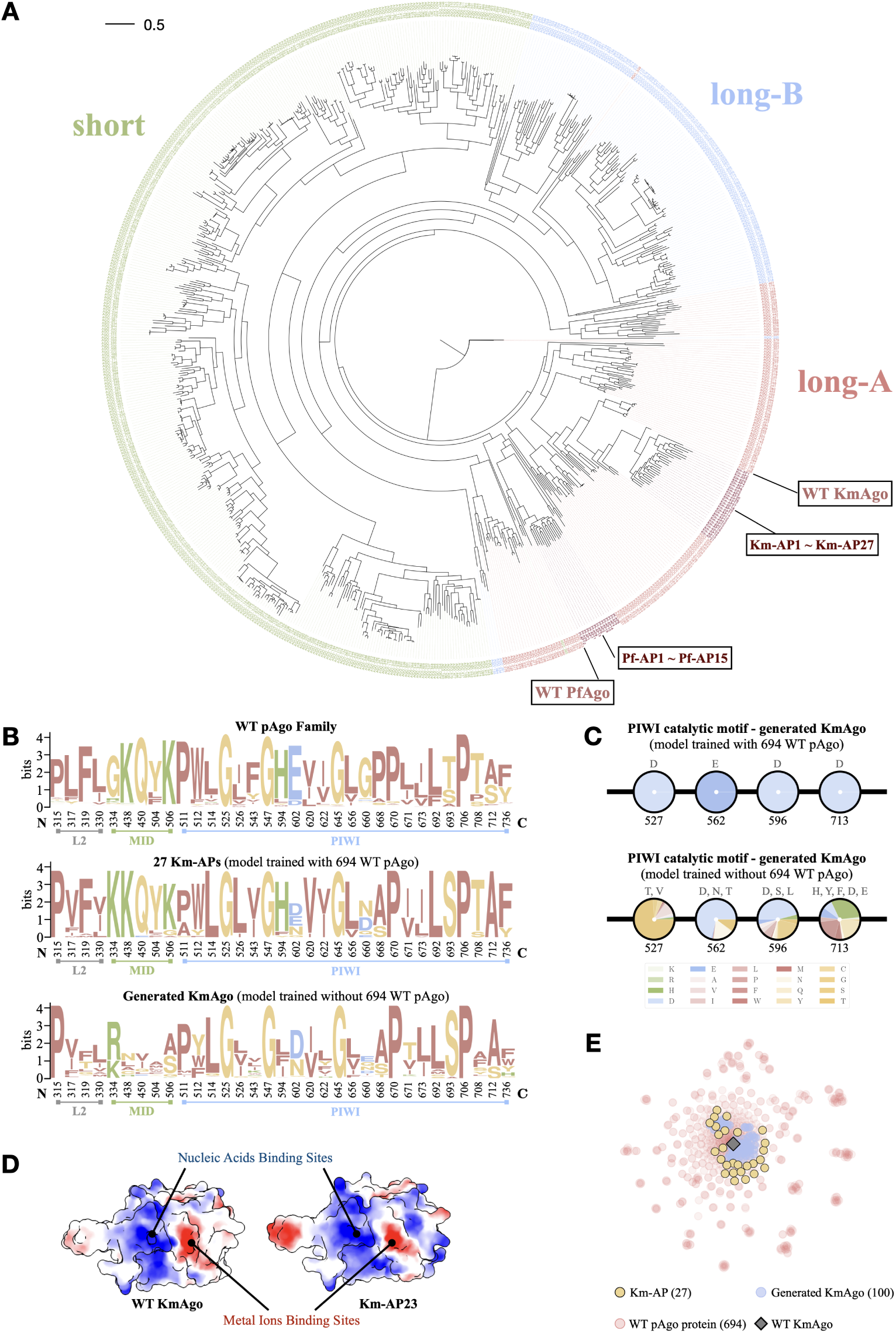
Evolutionary analysis on CPDiffusion-generated pAgo proteins. (**a**) Phylogenetic tree of natural pAgo protein sequences and generated (Km-APs and Pf-APs). (**b**) Conservative patterns of WT pAgo (upper panel), Km-APs (middle panel), and generated KmAgo sequences by CPDiffusion trained without the pAgo dataset (lower panel). The conservative positions are based on the alignment of the pAgo dataset. (**c**) AA composition at the catalytic motif (DEDD) for Km-APs (upper panel) and generated KmAgo sequences by CPDiffusion trained without the pAgo dataset (lower panel, detail in Fig. S100). Each pie chart denotes the AA composition of generated protein sequences at the site specified below the pie chart, and the top AA types with *>* 5% occurrence are listed above the pie chart. Training on the pAgo dataset significantly improves the model performance in preserving the conservative patterns for KmAgo. (**d**) Electrostatic surface of MID domain and PIWI domain in KmAgo and APs. Color ranges from red for negative potential through white to blue for positive potential. The arrows point out the nucleic acids binding sites and metal ions binding sites. (**e**) Computational Analysis of WT pAgo and generated KmAgos with t-distributed stochastic neighbor embedding (t-SNE). Each point represents a protein sequence embedding in a two-dimensional space for visualization purposes. The generated Km-APs by CPDiffusion tend to move from WT KmAgo toward the entire landscape of WT pAgo proteins.

Overall, the enhanced biofunctions observed in experimented Km-APs and Pf-APs, especially in terms of DNA cleavage activity, highlight the promising ability of CPDiffusion in function-oriented protein design. The model efficiently and effectively learns, discovers, and explores the implicit ‘sequence-structure-function’ relationship from a high-quality training dataset comprising a small set of proteins from the same family. This capability leads to the reliable generation of protein sequences with the desired functionality and enhanced performance.

### Bioinformatical and Computational Analysis of CPDiffusion-Generated Novel Sequences

To understand how the CPDiffusion-generated sequences relate to WT pAgo proteins in terms of their function and evolution relationship, we performed a comprehensive analysis based on their sequences and structures. First, we performed an evolutionary analysis on the Km-APs and Pf-APs to determine their relationship with other WT pAgo proteins. A phylogenetic tree was constructed through multiple sequence alignment (MSA) of Km-APs, Pf-APs, and proteins in the pAgo database (Fig. 5a, Methods). The designed Km-APs and Pf-APs, positioned within the long-A clade, belong to the KmAgo lineage and the PfAgo lineage, respectively, and share evolutionary properties with long-A pAgo proteins. We then delve into the conservative patterns captured by CPDiffusion-generated sequences at the 33 align-based conservative AA sites. For both KmAgo (Fig. 5b) and PfAgo (Supplementary Fig. S96), their generated sequences closely mirror the distribution of conserved AAs in the dataset. Additionally, the catalytic motif DEDD (for KmAgo, in Fig. 5c and Supplementary Fig. S97) and DEDH (for PfAgo, in Supplementary Fig. S98) are preserved perfectly in the generated sequences, the correct generation of which are pivotal for enabling pAgo proteins to achieve their catalytic functions [45]. We attribute the success of automatically capturing these crucial residues to the abundance of the high-quality pAgo dataset used to train the model. As supporting evidence, the generated sequences failed on both the DEDX motif and the conservative sites for recovering desired groups of AAs when the model was trained without pAgo proteins (bottom panels in Fig. 5b and c). Furthermore, the electrostatic surface properties of Km-APs and Pf-APs exhibit resemblances to those of the WT KmAgo and PfAgo, respectively, including the negatively charged core for metal ion binding at the cleavage site and the positively charged surface responsible for target nucleic acid binding (Fig. 5d, Supplementary Figs. S43 and S85). These findings affirm CPDiffusion’s ability to assimilate intrinsic sequential and structural characteristics, thereby ensuring the functional efficacy of the resultant proteins.

The observation of discernible variations in protein enzymatic activity, despite the overall similarity in APs’ structure to the WT’s, suggests the presence of distinct inter-residue interactions within the APs. To explore this, we selected six Km-APs (Km-AP8, Km-AP9, and Km-AP19 with reduced cleavage activity; Km-AP22, Km-AP23, and Km-AP27 with enhanced cleavage activity) and analyzed the impact of interactions in catalytic sites on the alteration in cleavage activity. The catalytic tetrad (D527, E562, D596, and D713) of the KmAgo protein forms a specific conformation to cleave the target DNA [38, 45]. In the case of Km-AP8, the E562 in the loop region of KmAgo has formed a small alpha-helix (highlighted in red in Supplementary Fig. S65), and this structural change might hinder E562 from inserting into the catalytic pocket due to increased steric hindrance. Regarding Km-AP9 and Km-AP19, there are missing beta-sheets near D527 (highlighted in orange and green in Supplementary Fig. S65), leading to a reduction in electrostatic interactions (Supplementary Table S18) and potentially destabilizing the structure of the catalytic pocket. Conversely, in the case of Km-AP22, Km-AP23, and Km-AP27, which exhibit higher activity than Km-WT, certain AAs within the turn fold into beta-sheets (highlighted in green in Supplementary Fig. S66). This observation indicates the presence of more hydrogen bonds and salt bridges compared to the WT (Supplementary Table S18). The increased interaction around the catalytic sites could enhance the structural stability of the catalytic pockets.

We further calculated the binding free energy of gDNA/tDNA in the catalytic pockets of Km-WT, Km-AP23 (enhanced function) and Km-AP9 (reduced function) using MD simulation (the structures of the protein-gDNA-tDNA complex are obtained from AlphaFold3). As shown in Supplementary Fig. S67, the binding free energy of gDNA/tDNA in the catalytic pocket of Km-AP23 is lower than that in Km-WT, while in Km-AP9 it is higher. This suggests that the binding affinity of gDNA/tDNA in the catalytic pocket of Km-AP23 is stronger, whereas in Km-AP9 it is lower compared to Km-WT. This stable binding state might facilitate the catalytic site’s cleavage of tDNA because the conformation of the catalytic pocket is crucial for the catalytic motif to cleave the target DNA [38, 45].

## 3 Discussions and Conclusions

This study introduces CPDiffusion, a novel pipeline for generating functional sequences tailored to a given protein backbone. We showcased the remarkable potential of this generative pipeline through its application to generate mesophilic endonucleases based on WT KmAgo and PfAgo with enhanced functionality. The generated novel sequences of KmAgo and PfAgo are less than 70% similar to their WT template sequences. For both groups of designed pAgo proteins, more than 90% novel sequences acquired DNA cleavage activity with over 70% observed activity enhancement over their WT baselines. Notably, the best-performing novel KmAgo achieves nine times better activity than WT KmAgo. The best novel PfAgo shifts the melting temperature of WT PfAgo from approximately 100 *^◦^*C to 50 *^◦^*C, and its ssDNA cleavage activity at 45 *^◦^*C is two times of that of the WT PfAgo at 95 *^◦^*C, which is 11 times over that of the WT KmAgo at moderate moderate temperature. These remarkable results demonstrate CPDiffusion’s strong potential in automatically learning from WT functional proteins and designing valid protein sequences with highly complex biofunctions toward enhanced functions.

Additionally, APs with enhanced functions could be potentially applied in *in vivo* biotechnology [25], particularly for cellular-level nucleic acid labeling. Their strong substrate affinity, stability within protein-substrate complexes, and precise targeting offer substantial advantages over conventional nucleic acid binding methods like DNA-painting and fluorescence in situ hybridization (FISH) [27, 46]. For instance, APs could assist guide DNA in binding to nucleic acids within mammalian cells (such as HEK293T and HeLa cells and fly embryos). The guide DNA carries overhang initiator sequences to initiate hybridization chain reactions (HCR), whereby multiple fluorophore-tagged secondary probes are recruited and assembled into a chain to amplify the signal. Compared to traditional FISH, pAgo-based FISH could stain nucleic acids within cells effectively and accurately at moderate temperatures without formamide, which is known to be toxic and a potential teratogen. Additionally, the high cleavage activity of APs could be leveraged for the specific cutting of labeled nucleic acids in cells, preparing the way for multiple rounds of nucleic acid labeling. Furthermore, the joint action of pAgo with enhanced function and nuclease-deficient RecBC helicase holds the potential to cleave double-stranded DNA [29], potentially leading to gene therapy approaches that precisely target genes responsible for diseases. Thus, the applications of APs could open up novel opportunities for developing therapies, *in vivo* imaging, cancer immunotherapy, and gene editing.

There has been considerable progress in designing new functional proteins using deep learning methods with experimental validations. ProteinGAN [6] learns the evolutionary relationships of protein sequences directly from the sophisticated multi-dimensional space of AA sequences. Out of 55 artificial sequences containing 321 AAs and one functional domain, 24% of the new proteins are soluble and display malate dehydrogenase catalytic activity. Pre-trained ProGen [8], a large language model trained on 280 million protein sequences and billions of network parameters, allows for conditional protein design. Through fine-tuning the model on a particular protein family, it generates lysozyme sequences (∼ 120 AAs) with low sequence identities, and thousands of protein samples exhibit activity. Lately, Watson et al. [9] introduced RFdiffusion based on structures, a function-oriented protein design method that achieves outstanding performance in protein monomer design, protein binder design, symmetric oligomer design, enzyme active site scaffolding, and symmetric motif scaffolding.

In comparison, CPDiffusion establishes a discrete denoising diffusion model for protein sequence generation characterized by protein-specific conditions, such as template secondary and tertiary structure, and extremely conserved AAs. Taking advantage of these specifications on the studied protein, valid sequences could be generated automatically by CPDiffusion with 4 million parameters trained on approximately 20, 000 family-diverse and 700 family-specific WT proteins. The model comprehends the protein’s intrinsic sequence-structure-function construction rules to guide sequence generation even for excessively long protein sequences (approximately 800 AAs) and highly complex functionality (six functional domains). CPDiffusion effectively learns the conservative patterns and other construction rules from the trained pAgo protein family. Remarkably, the pAgo sequences generated by CPDiffusion disperse from the template WT protein towards the landscape of the pAgo proteins family, encompassing diverse AA combinations and enzymatic mechanisms. Despite significant sequence differences, the proteins generated by CPDiffusion high similarity to natural proteins in the high-dimensional space. In other words, our method acquires knowledge of protein sequences, structures, and functionality in nature, facilitating exploration in latent space to unveil novel protein sequences with desired properties, such as thermostability, affinity (K*_M_*) and catalytic efficiency (k*_cat_*) to substrate, and overall activity (k*_cat_*/K*_M_*). In contrast to rational design or directed evolution, our deep learning-based protein generation strategy fosters a more extensive realm for exploring protein sequences, all while preserving functional integrity. Moreover, since our method exhibits a high success rate (*>* 90%) in generating diverse protein sequences (altering up to 400 AAs in a single step) with enzymatic activity, it offers a new avenue for protein engineering and the potential expansion of the sequence library and protein fitness landscape, providing researchers with a broader range of options to explore for desired protein functions within the family.

CPDiffusion is dedicated to generating novel AA sequences that exhibit low sequence homology with WT proteins, while demonstrating significantly improved enzymatic activity that better aligns with certain application requirements. While WT proteins, after hundreds of millions of years of natural evolution in distinct physiological conditions, may exhibit similar enzymatic functions with largely different sequences, the likelihood of failure is high when attempting to manually simulate this evolutionary process. CPDiffusion learns from a small set of training datasets, enabling the alteration of hundreds of AAs in a single step to design novel multi-domain sequences. Remarkably, CPDiffusion achieves a success rate of over 90% and successfully discovers highly active sequences that have not been reported in the literature. The underlying generation strategy also plays a pivotal role in expanding the available reservoir of potential protein candidates, thereby establishing a paradigm shift in the exploration of existing proteins. Additionally, our end-to-end generation method facilitates protein engineering endeavors. When engineering a protein toward desired functionalities, our method does not require time-consuming and costly iterative selections like directed evolution methods, which start from single-site mutations and gradually combine multiple mutations through lengthy iterative processes. Meanwhile, the progressive denoising steps integrated within our CPDiffusion can be perceived as guided pathways leading from arbitrary initial sequences toward their optimized states in a latent space. Our model streamlines the generation and evolution process by enabling the direct creation of proteins that manifest commendable biological performance in a single step. This approach, in turn, presents a myriad of innovative starting points that propel directed evolution toward the swift discovery of novel sequences possessing superior properties. The efficiency and efficacy exhibited by CPDiffusion instill promising prospects for expediting the protein design and discovery.

## 4 Methods

### 4.1 Conditional Protein Denoising Diffusion

DDPMs [11] approximate a distribution by parameterizing the reversal of a discrete-time forward diffusion process. In the context of protein sequence generation, the forward diffusion operates on the distribution of AA types 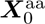 that involves the iterative noise addition for *T* steps until reaching 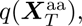 a distribution indistinguishable from a reference independent transition distribution. The forward diffusion 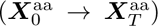 samples 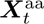 at every step *t* (*t* = 1*, . . ., T*) using the forward diffusion kernel defined as 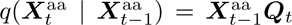 until 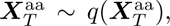 where ***Q****_t_* denotes the transition probabilities at *t*. The reverse diffusion 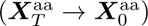 samples 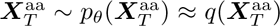 to convert from the prior into samples of the learned data distribution, *i*.*e*., 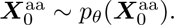 During the progressive training process, the generative probability distribution 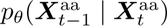 is parameterized by a neural network *f* (*θ*) that embeds additional characterization of protein instances, such as their topology and physicochemical features. We represent proteins as graphs with nodes corresponding to AAs and implement equivariant graph neural networks (*e*.*g*.,, EGNN [47]) that perform rotation and translation equivariance on aggregating node attributes based on relative spatial relationships of corresponding neighboring AAs. Additional conditions are also possible to be inserted to guide the denoising process, such as the secondary structure and transition matrix. Notably, we defined a conserved AA as the known region and conditioned the reverse diffusion process of an unconditional DDPM for sequence inpainting [14]. We fixed D713 for the WT KmAgo [32, 43], which is the position of the catalytic site that depends on the particular Ago of interest. The weights in the neural network *f* (*θ*) are optimized by minimizing the cross-entropy ℒ_CE_ between 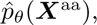 the network prediction that approximates *p_θ_*(***X***^aa^), and the observed AA types. Further details regarding the diffusion model can be found in SI.

### 4.2 Structure Prediction and Comparison

To predict the structure for the artificial Ago sequences generated by conditional protein denoising diffusion, we used AlphaFold2 in single-sequence mode with multiple sequence alignment (MSA) and with PDB templates. The highest-ranked predicted structure among the five models was used. The RMSD between WT and artificial Ago protein in our study are calculated using PyMOL (The PyMOL Molecular Graphics System, Version 2.0, Schrödinger, LLC.). Hydrogen bonds are defined as polar contacts in proteins, salt bridges are defined by locating oxygen atoms in AAs, and nitrogen in base AA pairs has distances within 4 Å.

### 4.3 Prokaryotic Argonaute Protein Database

We constructed a dataset containing 694 pAgo proteins, selected from a comprehensive Ago dataset [37]. This curated set represents a broad diversity of pAgo types, including short, long-A, and long-B pAgo proteins. We performed all five models that AlphaFold2 [39] provides to predict the structures of WT pAgo sequences. Since the results using five different models are generally consistent, we used the results of all proteins via Model 1 for further analysis. In Supplementary Figs. S100-S101, we delve into the diversity of the pAgo database to alleviate concerns regarding data bias and highlight its crucial role in enhancing the performance of CPDiffusion. Generally, pAgo proteins in the dataset exhibit structural conservation alongside sequence diversity. Furthermore, the majority of WT proteins are mesophilic pAgos, with a few long-A pAgo proteins belonging to the thermophilic category.

### 4.4 Sequence and Structure Analysis

The model-generated Ago proteins and 694 Ago proteins from the database were subjected to multiple sequence alignment using the MUSCLE v5 software with the super5 algorithm [48]. Conservation scores for each residue were computed by

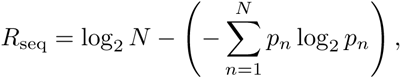

where *p_n_* is the observed frequency of AA *n*, and N=20 for all 20 kinds of AAs. Conservation score equals *R*_seq_ multiplied by the fraction of non-gapped positions in the column to apply the gap penalty. Residues in the Ago database scoring above 2.5 were selected. Residues in the design scoring below 0.2 were excluded to mitigate alignment discrepancies (Supplementary Fig. S99). The conserved residues were visualized using WebLogo [49]. The phylogenetic tree is computed via IQ-TREE v1.6 with 1500 ultrafast bootstraps and used BLOSUM62 as a substitution model [50]. FigTree v1.4.4 is used to visualize the phylogenetic tree (http://tree.bio.ed.ac.uk/software/figtree/). The electrostatic surface, or the Coulombic electrostatic potential (ESP) of the protein was subsequently assessed using the ‘coulombic‘ command in Chimera visualization tool, which provided a graphical display of the distribution of electrostatic potential on the surface of the protein.

### 4.5 Protein Expression and Purification in BL21(DE3)

A codon-optimized version of KmAgo, Km-APs, PfAgo, Pf-APs, BlAgo, PbAgo, CbAgo, and SeAgo genes was synthesized by Sangon Biotech (Shanghai, China). It was cloned into the pET15(b) plasmid (the construction of Ago protein is shown in Supplementary Figs. S48-S49) to construct pEX-Ago with an N terminal His-tag. The expression plasmid was transformed into *Escherichia coli* BL21(DE3) cells. A 30 ml seed culture was grown at 37 *^◦^*C in LB medium with 50 *µ*g/ml kanamycin and was subsequently transferred to 500 mL of LB in a shaker flask containing 50 *µ*g/ml kanamycin. The cultures were incubated at 37 *^◦^*C until the OD600 reached 0.6-0.8, and protein expression was then induced by the addition of isopropyl-D-thiogalactopyranoside (IPTG) to a final concentration of 0.5 mM, followed by incubation for 16-20 h at 18 *^◦^*C. Cells were harvested by centrifugation for 30 min at 4,000 rpm, and the cell pellets were collected for later purification. The cell pellets were resuspended in lysis buffer (25 mM Tris-HCl, 500 mM NaCl, 10 mM imidazole, pH 7.4) and then disrupted using a High-Pressure Homogenizer at 700-800 bar for 5 minutes (Gefran, Italy). The lysates were centrifuged for 30 min at 12,000 rpm at 4 °C, after which the supernatants were subjected to Ni-NTA affinity purification with elution buffer (25 mM Tris-HCl, 500 mM NaCl, 250 mM imidazole, pH 7.4). Further gel filtration purification using a Superdex 200 (GE Tech, USA) was carried out with an elution buffer. The fractions resulting from gel filtration were analyzed by SDS-PAGE. The fractions fractions containing the protein were flash frozen at -80 *^◦^*C in buffer (15 mM Tris–HCl pH 7.4, 200 mM NaCl, 10% glycerol).

### 4.6 Single-Strand DNA/RNA Cleavage Assay

For standard activity assays of pAgo proteins and APs, cleavage experiments were performed in a 2:1:1, 5:2:2, or 3:2:2 molar ratio (protein:guide:target). For studying the effect of temperature on the activity of proteins, they were incubated at 42 *^◦^*C for 2 min and 5 min before conducting the ssDNA cleavage assay. First, 5 *µ*M protein was mixed with a synthetic 1 *µ*M gDNA guide in the reaction buffer (15 mM Tris-HCl (pH 7.4), 200 mM NaCl, and 5 mM MnCl_2_). The solution was then pre-incubated at 37 *^◦^*C for 20 min. After pre-incubation, 1 *µ*M tDNA, which was labeled with the fluorescent group 6-FAM at the 5′-end and the quencher BHQ1 at the 3′-end, was added to the mixture. The fluorescence signals were traced by the quantitative real-time PCR QuantStudio 5 (Thermo Fisher Scientific, USA) with *λ*_ex_ = 495nm and *λ*_em_ = 520nm. The results were analyzed by QuantStudioTM Design & Analysis Software v1.5.1. The guide and target nucleic acids used for cleavage are listed in Supplementary Table S16 and S17. Two independent experiments are conducted to determine the cleavage activity of Ago proteins. The original data is shown in Supplementary Figs. S55-S59 and S92. For the kinetic measurements, the parameters k*_cat_* and K*_M_* were determined by fitting the Michaelis–Menten equation to the velocity of each reaction as a function of the concentration of ssDNA target.

### 4.7 Circular Dichroism Spectroscopy

The CD measurements on the secondary structure of WT and APs were performed in a Jasco J-1500 spectropolarimeter with a 1 mm pathlength cell. The concentrations of proteins were 0.1 mg/mL in 1× PBS buffer (pH = 7.4). For CD spectra, we used PBS buffer as the solvent instead of Tris-HCl buffer because the Tris-HCl buffer has a strong CD background, which will significantly affect the analysis of the CD signal of proteins [44]. The original data is shown in Supplementary Fig. S51.

### 4.8 Fluorescence Polarization Assay

To determine the apparent dissociation constant (Kd) for proteins binding to either gDNA or tDNA, a fluorescence polarization assay was conducted using a multifunctional enzyme-linked immunosorbent assay plate reader (Spark, Tecan). A solution was prepared by combining 5 nM of 3’ 6-FAM labeled guide or target DNA with proteins across a concentration range of 0 to 1500 nM in a reaction buffer (15 mM Tris-HCl at pH 7.4, 200 mM NaCl, and 5 mM MnCl_2_). This mixture was incubated at 37°C for 1 hour and subsequently transferred to a light-protected 96-well ELISA plate. The degree of polarization was measured using the Spark Tecan plate reader, employing an excitation wavelength of 485 nm and an emission wavelength of 525 nm. All experiments were independently conducted three times. The binding percentages were analyzed using Microsoft Excel and Prism 8 (GraphPad) software. The data was fitted with the Hill equation, incorporating a Hill coefficient of 2 to 2.5. The original data is shown in Supplementary Fig. S54.

### 4.9 Cell-Free Protein Expression and Extraction

The expression of KmAgo and APs was accomplished utilizing the Tierra Bioscience cell-free expression platform. The preparation of cell-free extracts for protein expression was conducted in adherence to the methodology established by Sun et al [51].

### 4.10 Differential Scanning Fluorimetry

Each protein sample (PfAgo and Pf-APs) containing 1 *µ*M of protein in a buffer containing 15 mM Tris-HCl (pH = 7.4) and 200 mM NaCl was prepared in triplicate and added to PCR tubes. SYPRO Orange dye available as 5000× stock (Sigma-Aldrich) was added just before the measurement of proteins in an appropriate amount to achieve a final concentration of the dye of 5×. The thermal denaturation of proteins was monitored by exciting the SYPRO Orange dye at 470 nm and monitoring its fluorescence emission at 570 nm using Q-PCR (Analytikjena, Germany). The baseline correction is used by the Opticon Monitor software available on the PCR instrument. The original data is shown in Supplementary Fig. S95.

### 4.11 Small Angle X-ray Scattering

Small angle X-ray scattering (SAXS) measurement was employed to the status of proteins in buffer. Synchrotron SAXS measurements were conducted at BL19U2 beamline in Shanghai Synchrotron Radiation Facility (SSRF). The X-ray wavelength was 0.103 nm. Protein samples were dissolved in a buffer containing 15 mM Tris-HCl (pH 7.4), 200 mM NaCl. The concentration of samples is 0.5 mg/ml. Protein solutions were loaded into the silica cell and then gently refreshed with a syringe pump to prevent X-ray damage. To calculate the absolute intensity of protein, the empty cell and buffer were also measured at corresponding temperatures. Two-dimensional (2D) diffraction patterns were collected by the Pilatus 2 M detector with a resolution of 1043 × 981 pixels of 172 *µ*m × 172 *µ*m. Twenty sequential 2D images were collected with 0.5 s exposure time per frame. The 2D scattering patterns were then integrated into one-dimensional (1D) intensity curves by using Fit2D software from the European Synchrotron Radiation Facility (ESRF). Frames with no radiation damage were used for further processing. The one-dimensional data is processed by using ScÅtter and ATSAS software.

### 4.12 Molecular Dynamics Simulations

The structures of proteins with nucleic acids for simulations were predicted from AlphaFold3. Protein and a large number of water molecules were filled in a cubic box. 16 chlorine counter ions were added to keep the system neutral in charge. The CHARMM36m force field was used for the complex and the CHARMM-modified TIP3P model was chosen for water. The simulations were carried out at 310 K. After the 4000-step energy-minimization procedure, the systems were heated and equilibrated for 100 ps in the NVT ensemble and 500 ps in the NPT ensemble. The 100-ns production simulations were carried out at 1 atm with the proper periodic boundary condition, and the integration step was set to 2 fs. The covalent bonds with hydrogen atoms were constrained by the LINCS algorithm. Lennard-Jones interactions were truncated at 12 Å with a force-switching function from 10 to 12 Å. The electrostatic interactions were calculated using the particle mesh Ewald method with a cutoff of 12 Å on an approximately 1 Å grid with a fourth-order spline. The temperature and pressure of the system are controlled by the velocity rescaling thermostat and the Parrinello-Rahman algorithm, respectively. All MD simulations were performed using GROMACS 2020.4 software packages.

### 4.13 Electrophoretic Mobility Shift Assay

To examine the loading of gDNA onto proteins, proteins and 3’ end FAM-labeled guide were incubated in 20 µL of reaction buffer containing 15 mM Tris-HCl pH 7.4, 200 mM NaCl for 20 min. The concentration of 3’ end FAM labeled guide and protein was fixed as 1 µM and 2 µM, respectively. Then the samples were mixed with 5 µL 5× loading buffer (Tris-HCl (pH 7.4), 25% glycerin, Bromophenol blue) and resolved by 12% native PAGE. Nucleic acids were visualized using Gel DocTM XR+.

## Supplementary Information

Supplementary Figs S1-S102, Supplementary Tables S1-S24.

## Author Contributions

B.Zhou and L.Z. conceived the concept, designed the experiments, and wrote the original manuscript. B.Zhou, K.Y., and Y.T. developed the CPDiffusion. L.Z., B.W., and Q.L. performed the wet lab experiments. B.Zhong conducted the bioinformatics analyses. B.Zhou, L.Z., P.L., and L.H. revised the manuscript.

## Acknowledgments

The authors acknowledge the Center for High-Performance Computing at Shanghai Jiao Tong University for computing resources.

## Additional Information

Correspondence and requests for materials should be addressed to, and.

## Data availability

All the data are presented in Maintext and SI.

## Code availability

The model implementation can be found at https://github.com/bzho3923/CPDiffusion.

